# Evaluation of Iodine-Based Contrast Agents for Ex Vivo Micro-CT Imaging of Porcine Hearts: An Application to Cardiac Conduction System Imaging

**DOI:** 10.64898/2025.12.12.693377

**Authors:** Manu Pradeep, Shuvashis Das Gupta, Yi Li, Sami Kauppinen, Mikko Finnilä, Timo Liimatainen

**Affiliations:** Research Unit of Health Sciences and Technology, University of Oulu, Oulu, Finland; Department of Biomedical Engineering, Lund University, Lund, Sweden; Biocenter Oulu, University of Oulu, Oulu, Finland; Department of Diagnostic Radiology, Oulu University Hospital, Oulu, Finland

**Keywords:** Cardiac conduction system, SAN, AVN, Purkinje fiber, Contrast-enhanced micro-computed tomography, iodine

## Abstract

**Objective:** Visualization and segmentation of the cardiac conduction system (CCS) enable advanced mathematical modeling and improve the diagnosis of heart rhythm disorders.

**Background:** This study investigates iodine in ethanol (I_2_E) and Lugol’s solution (I_2_KI) for optimizing high-resolution micro-computed tomography (micro-CT) imaging of the CCS in porcine hearts. The study evaluates their efficacy in enhancing tissue contrast, shrinkage, and downstream processing.

**Methods:** Dissected porcine hearts were stained with I_2_E and I_2_KI for contrast enhancement and scanned with micro-CT system. Volumetric shrinkage of tissues during staining was evaluated. Additionally, qualitative visualization of CCS-related anatomical landmarks, such as the sinoatrial node (SAN), atrioventricular node (AVN), and Purkinje fibers, was performed, along with assessment of artifact occurrence and sample integrity. The regions corresponding to CCS were segmented from micro-CT image data and were then validated against histology.

**Results:** Although both contrast agents yielded comparable Signal-to-Background ratios (p = 0.8871), I_2_E demonstrated significantly superior imaging quality with a higher Contrast-to-Noise Ratio (p ≤ 0.05) and lower overall noise (p ≤ 0.05) than I_2_KI. Furthermore, while I_2_KI offered faster staining, it caused leaching, artifacts, and structural disintegration that compromised downstream applications. In contrast, I_2_E preserved sample integrity and minimized artifacts, facilitating seamless post-processing and histological sectioning.

**Conclusions:** These results suggest that I_2_E could be a viable alternative to I_2_KI for CCS micro-CT imaging when sample preservation and downstream analyses are essential.

## Introduction

Imaging has been a crucial segment in cardiovascular research, particularly that of the cardiac conduction system (CCS). When it comes to cardiac-related animal studies, porcine hearts are the most preferred choice since, anatomically and physiologically, they closely resemble the human heart, making them a valuable model for cardiac research [1,2]. Ex vivo animal research enables longer scan durations without concerns for anaesthesia or radiation exposure. Moreover, motion artifacts from breathing and heartbeat can be avoided in this setting.

Contrast-enhanced micro-computed tomography (micro-CT) has significantly improved visualization of cardiac tissues, but the lack of standardized protocols results in substantial tissue shrinkage, morphological distortion, and artifacts [3]. Therefore, an optimized micro-CT-based protocol for 3D imaging of the major regions of the CCS with minimal sample damage and reversible contrast enhancement for histopathological assessment in relatively large mammalian hearts remains to be developed.

The CCS is responsible for transmitting electrical impulses that, in turn, synchronize the periodic contractions of the atria and ventricles of the heart. The CCS consists of four anatomical structures: the sinoatrial node (SAN), the atrioventricular node (AVN), the bundle of His, and the Purkinje fibers (PF). Following these structures, the electrical impulse is transmitted to the myocardium, ultimately leading to myocardial contraction and conduction of the signal throughout the heart [4]. Previous studies have imaged CCS structures both in vivo and ex vivo, but poor spatial resolution and safety concerns in diagnostic modalities remain bottlenecks for research [5–8]. In such scenarios, preclinical imaging modalities such as micro-CT can be used, providing significantly higher spatial resolution in 3D [9].

I_2_KI has been the most preferred staining agent in cardiovascular micro-CT imaging, although there have been few studies with other agents, such as phosphotungstic acid (PTA), graphene oxide, and osmium tetroxide[10– 13]. PTA has been reported to have poorer contrast, a highly acidic nature that causes shrinkage, and a slower rate of diffusion when compared to iodine-based contrast agents [14]. Other heavy-metal-based agents, such as osmium tetroxide, are toxic and require careful handling. Even though I_2_KI is non-toxic, it has been found to pose a bottleneck during post-imaging and histological sectioning, compromising sample integrity.

The use of iodine in ethanol (I_2_E) as a contrast agent has been explored in various soft-tissue micro-CT imaging studies. Similar comparative studies between contrast agents (PTA, Gadolinium) have been conducted, and it has been reported that I_2_E staining provides higher contrast in avian cephalic specimens and ex vivo lung imaging[15,16]. Another study comparing I_2_E and I_2_KI reported that I_2_KI-stained images appeared grainier and indicated a higher acidity level[17].

This exploratory study assesses whether I_2_E could serve as a viable alternative as a contrast agent for ex vivo micro-CT imaging of thick porcine cardiac tissues. It aims to provide a preliminary imaging pipeline for I_2_E and I_2_KI by comparing initial observations on their staining behaviour, structural preservation, and downstream histological integrity.

## Methods

### Preparation of samples

A total of 10 cardiac sections (2 each of SAN, AVN, apical region containing Purkinje fibers, and 4 myocardial sections) were dissected from two porcine hearts obtained from the local slaughterhouse. One set of samples (n_1_=4) underwent I_2_E staining, and the other set (n_2_=4) underwent I_2_KI staining. Additionally, the remaining two myocardial sections were utilized to evaluate shrinkage for each contrast agent. The dissected samples are represented in Fig 1. All the samples were first fixed in formalin and stored at 4 degrees Celsius.

**Fig 1.**
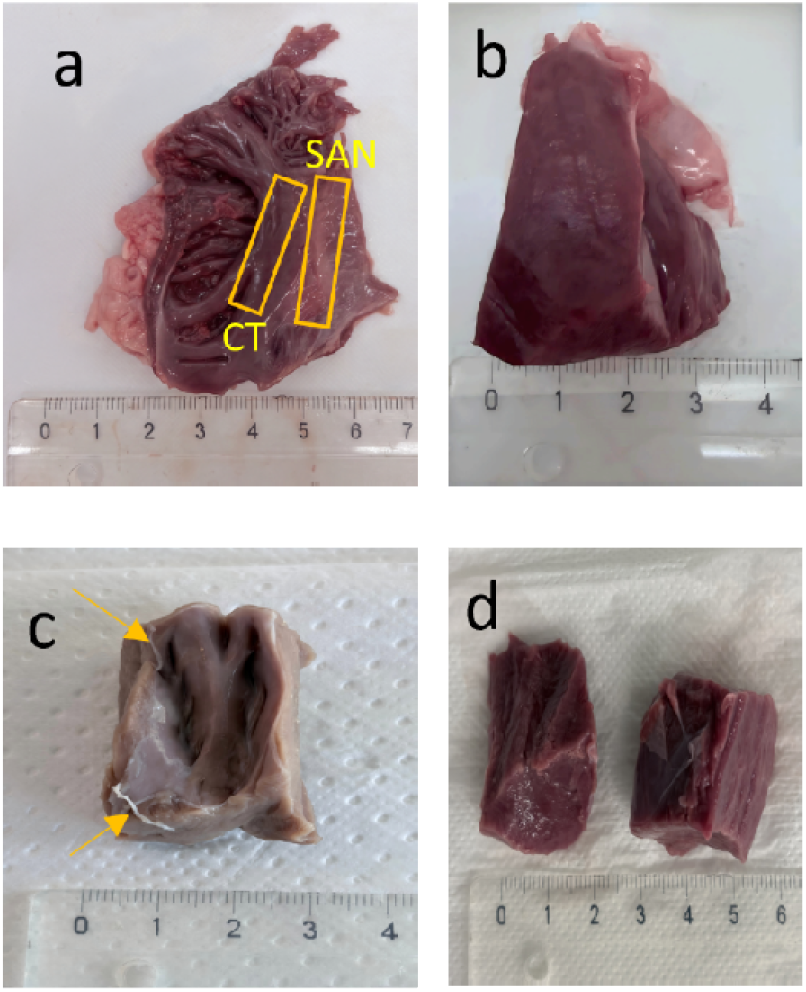
(a) Dissected SAN region from the porcine heart; CT: Crista Terminalis, (b) AVN section, (c) Formalin-fixed apex section (arrows indicate the larger false tendons), (d) Myocardium sections

### Preparation of chemicals

A 2.5% I_2_KI solution was prepared in distilled water. Previous studies have taken the concentration of I_2_KI to vary mainly from 1% to 10%[3,18,19]. However, owing to the acidification of I_2_KI over time, a concentration of 2.5% is adopted in this study, and the protocol was followed as reported in the study by Simcock et al [19]. A 1% iodine solution in ethanol (I_2_E) was also prepared, and the solutions were magnetically stirred for 30 minutes, then wrapped using aluminium foil, and stored in a dark environment. To stabilize the samples for micro-CT imaging, agarose was used as the embedding medium. For optimization and quantification purposes, smaller myocardium tissue sections were used, which could be stabilized by 1% agarose in PBS. However, for the larger CCS sections, a 3% agarose solution in PBS was prepared. The solution was microwaved for around 3 minutes to dissolve the agarose completely. To attain the gelling temperature, the solution was heated in a water bath at 37 degrees Celsius.

### Micro-CT Imaging

The dissected cardiac sections were used to perform the two workflows (Fig 2). Initially, the formalin was removed from samples by placing them in running tap water overnight. For the I_2_E approach, the sample required a dehydration procedure before staining. To achieve this, the sample underwent dehydration with ascending ethanol washes (30%, 50%, 70%, 80%, 90%, 100%, 100%). Each wash lasted 3 hours for the CCS structures since the structures were thick, and the smaller myocardium samples were subjected to 30 minutes of dehydration. To quantify shrinkage, the myocardium samples were used since they are smaller in size and easy to prepare. It was necessary to perform a scan on unstained myocardium so that their native volume could be obtained. These sections were exclusively used for shrinkage analysis and only their outer tissue boundary was required to calculate the volume. For this, the first myocardium sample was scanned before and after dehydration, and the second myocardium sample was scanned immediately after formalin removal. These were optimized to be quick scans of under ten minutes. The first day of staining began after these native scans. The section was then immersed in the prepared I_2_E solution. For the I_2_KI approach, another section was directly immersed in the prepared I_2_KI for the same periods. The stained samples were then fixed in 1% agarose and subjected to micro-CT imaging every 24 hours until day 5 of staining, using the Zeiss XRadia Versa 610 scanner. Unstained myocardium samples were scanned using consistent parameters: 40 kV voltage, 3 W power, 1.3 s exposure time, and a binning of 8. The pixel sizes were between 80-120 microns.

**Fig 2.**
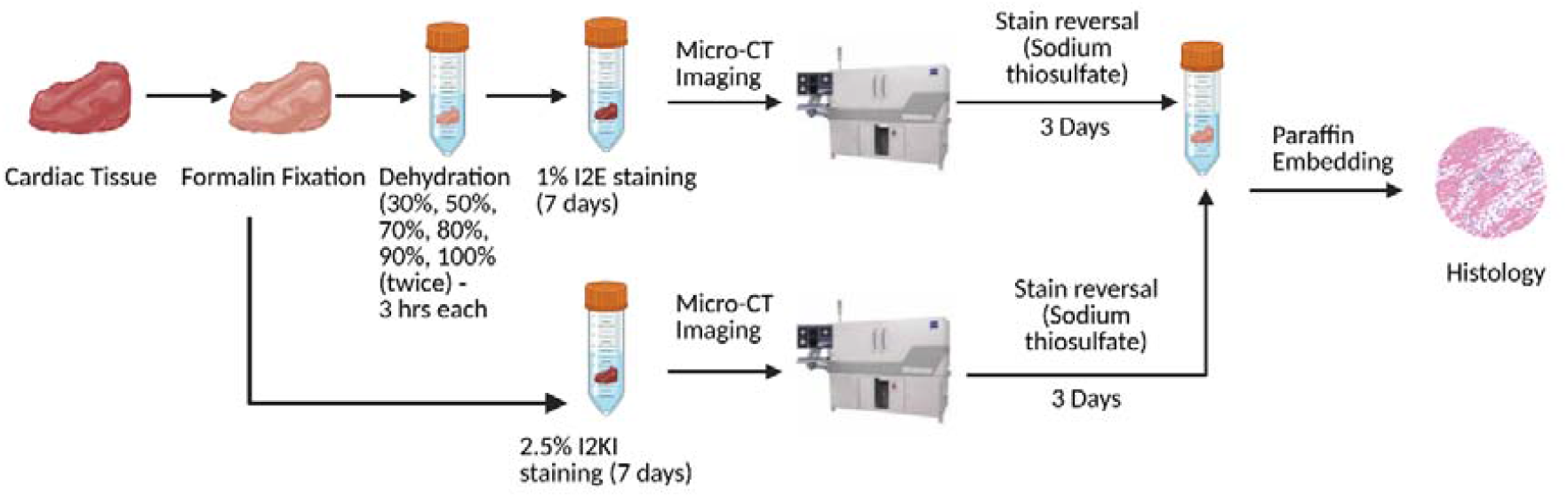
Workflow of the imaging protocol for both I_2_E and I_2_KI-stained samples (Created in https://BioRender.com)

The same protocols were followed in imaging the CCS structures as well. Since the samples were considerably larger than the myocardium, 7 days of staining was done for both contrast agents. Agarose embedding prevented the sample from drying during more extended scan periods and provided sufficient background separation in the images. The imaging using consistent settings: 60 kV voltage, 6.5 W power, 2 s exposure time, a Zeiss Low-Energy filter 6, and a binning of 2. The pixel sizes were between 35-50 microns. The image data were then imported into the ORS Dragonfly software (Dragonfly 2022.2 [Computer software], Comet Technologies Canada Inc., Montreal, Canada) for image visualization and segmentation.

### Histological Examination

To reverse the iodine staining, a 3% sodium thiosulfate (STS) solution in 70% ethanol was prepared, and the samples were immersed in it for 3 days after micro-CT imaging. Following contrast agent removal using STS, samples were histologically processed and embedded in paraffin, and thin sections were stained with Masson’s trichrome for histological evaluation. Both histology and micro-CT images were qualitatively compared.

### Image Analysis

To analyse the image quality obtained with I_2_E and I_2_KI, micro-CT image data from the cardiac samples were utilized. The signal-to-background (S-B) and contrast-to-noise ratios (CNR) were calculated from the grey values of the reconstructed images. A fixed range of attenuation coefficients was used to map voxel values to greyscale, to have a consistent dynamic pixel value range, and to allow direct comparison between the samples. Image quality was analyzed with ImageJ by drawing rectangles inside the tissue and background (agarose). This directly gave the intensity values inside the rectangular regions [20,21]. The ratio of the mean greyscale values of both the tissue and background was taken to interpret the S-B ratio. The S-B ratio was thus calculated based on the formula given by:

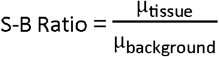

The CNR was calculated based on the formula given below:

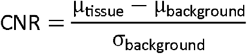

where μ is the mean grayscale value, and σ is the standard deviation.

The noise artefacts during micro-CT imaging were also separately calculated as the standard deviation (σ) of the background which would point to the leaching of contrast agents into the agarose.

The *p*-values (CI = 95%) were calculated via Welch’s t-test for S-B Ratio and CNR, and Mann-Whitney U test for Noise. Welch’s t-test, a robust parametric method, was specifically selected to evaluate the continuous S-B Ratio and CNR metrics to account for the unequal variances observed between the experimental groups. This approach minimizes the risk of false-positive errors within the limited sample cohort, ensuring that the calculated significance accurately reflects true differences in signal performance. The Mann-Whitney U test, a non-parametric method, was used to assess the Noise metric because of high variability and the existence of outliers in the limited sample group. This approach ensures that the detected statistical significance represents a true disparity in the noise distributions of the contrast agents, rather than an erroneous outcome influenced by an outlier measurement.

The shrinkage values for micro-CT images were quantified as the decrease in sample volume after each day of staining using the ORS Dragonfly software. For this, the sample was manually segmented out from the surrounding agarose, and the volume of this 3D ROI was calculated. For the CCS sections, the images were used to visualize and segment SAN, AVN, and Purkinje fibers using the same software.

## Results

The larger size of the CCS samples necessitated the use of a higher concentration of agarose for embedding. A low concentration of agarose (<3%) made the sample unstable during the scan, leading to image blurring. However, 1% agarose was found to be sufficient for imaging the smaller myocardial sections.

### Qualitative Analysis of the Cardiac Sections from micro-CT and histology images

From the micro-CT scans, samples stained with I_2_E showed better visual contrast than those stained with I_2_KI. Additionally, the I_2_KI-stained samples were observed to have cracks within the sample, compromising the sample integrity (Fig 3a,3b). There was also disintegration of the sample, observed both during and after the scans. The better contrast of I_2_E further helped in segmenting the SAN and arteries, the penetrating bundle (PB) that, in turn, bifurcates into left and right branches of the bundle of His, and part of the Purkinje fiber network (Figure 3c-g). Purkinje fibers appeared almost with the same contrast as that of the surrounding myocardium in the high-resolution micro-CT scan. These fibers were observed to be present within the false tendons and embedded within the extracellular matrix of the endocardium, having a diameter of roughly 60-100 microns (Fig 3f,3g). Even then, those fibers located at the ventricular walls were not identified due to insufficient contrast. The 2D and 3D (segmented) images are shown in Fig 3 and Fig 4, respectively.

**Fig 3.**
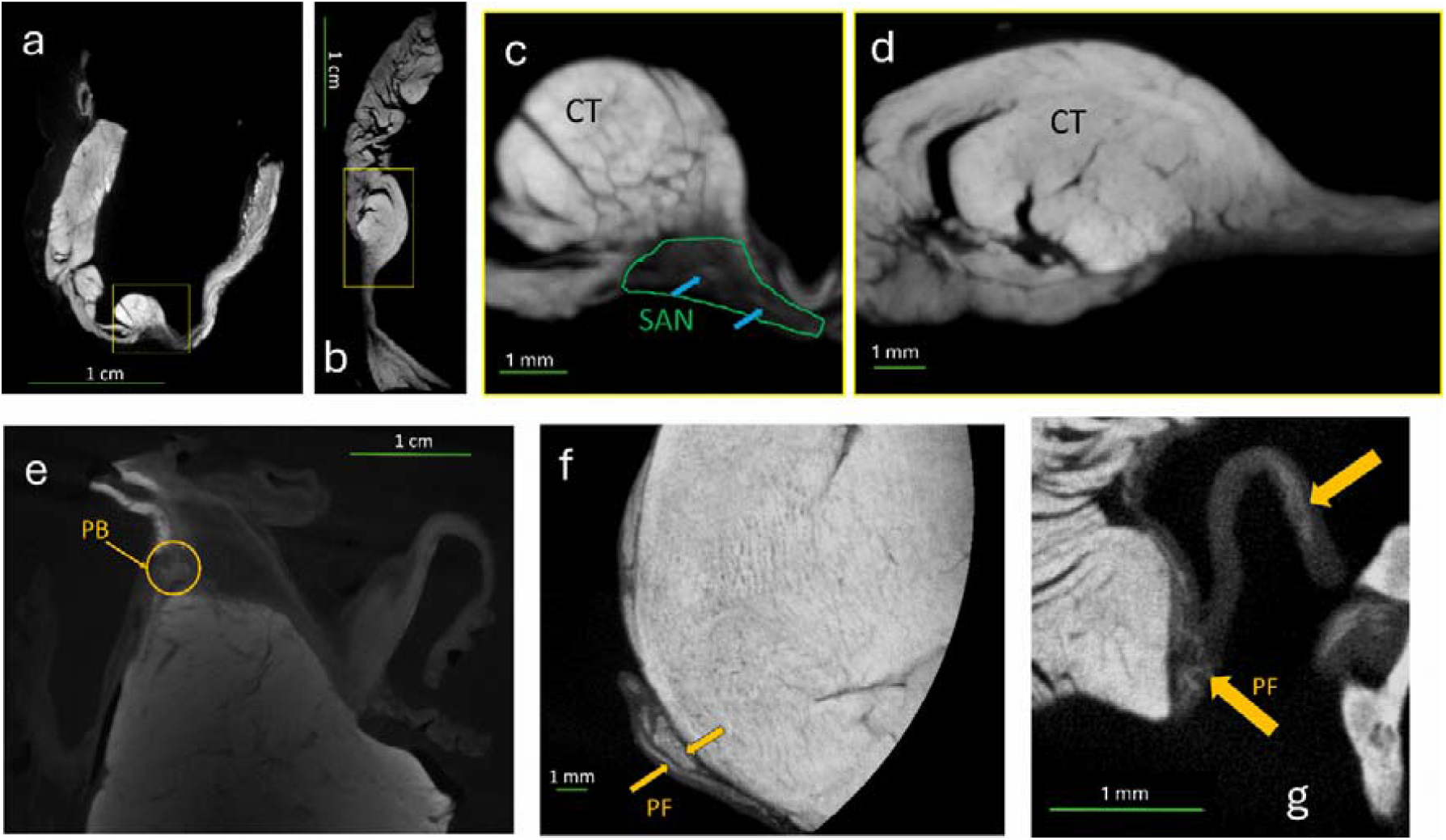
Micro-CT images of (**a**) I_2_E stained SAN section (**b**) I_2_KI stained SAN section, (**c**) (**d**) Magnified images of marked regions in (a) and (b) respectively, showing the crista terminalis (CT), SAN region and SAN arteries (indicated by the blue arrows). Compared to (a), the contrast in (b) is poor and does not clearly differentiate the anatomical landmarks (**e**) I_2_E stained AVN section showing the penetrating bundle (PB) (**f**) (**g**) Magnified images of the Purkinje fibers (PF-yellow arrows indicate the fibers running through the endocardium and false tendons respectively)

**Fig 4.**
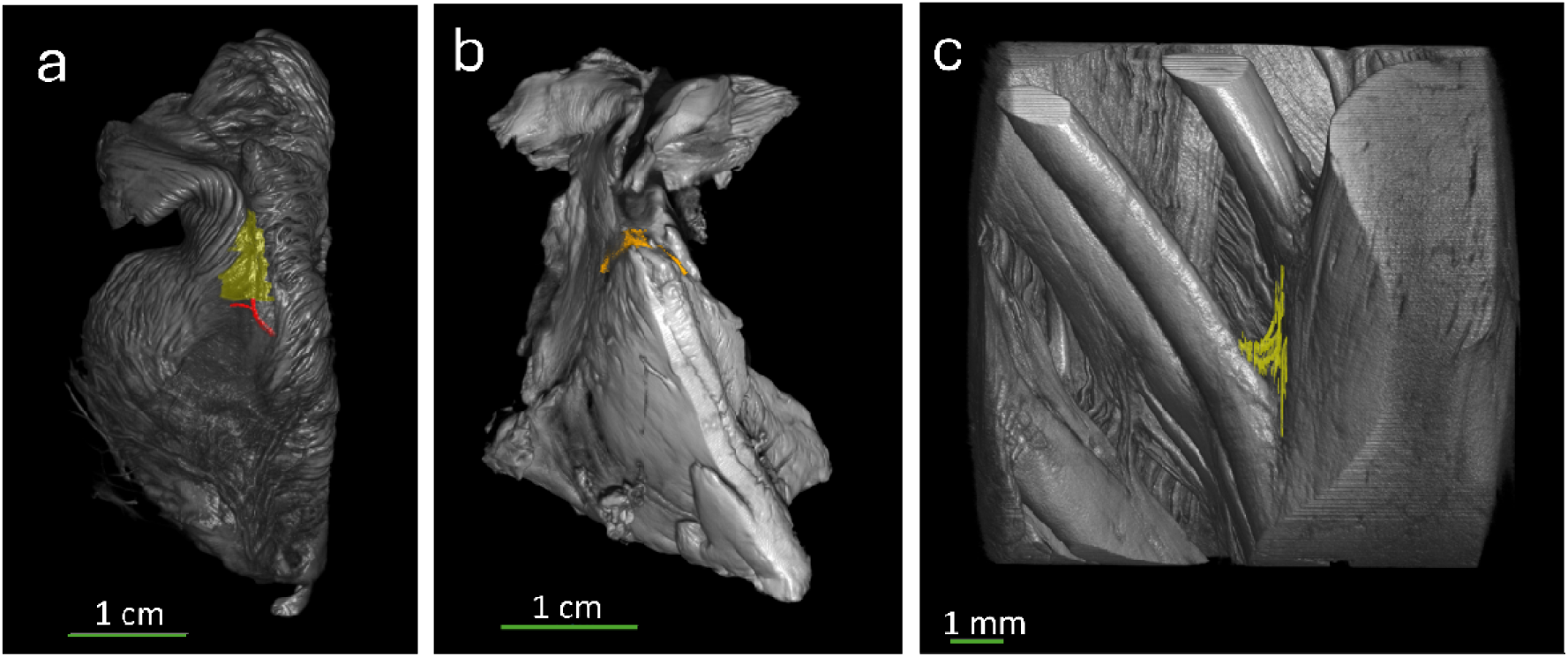
3D visualization of the CCS using micro-CT imaging (all images are from I_2_E stained sections, since I_2_KI offered poor anatomical delineation): **(a)** Segmentation of SAN (yellow) and SAN arteries (red); **(b)** the left and right bundle branches from the AVN section; **(c)** High resolution scan of apex region showing partial segmentation of Purkinje fibers due to poor contrast between the Purkinje fibres and False tendons

Upon comparison with histology sections, all landmarks were clearly observed in both I_2_E and I_2_KI-stained samples. The I_2_KI sections nevertheless showed higher disintegration in the histology sections (comparing Fig 5a, b and Fig 5c,d). The SAN was identified along with the bifurcating arteries (Fig 5c,d). The AVN section showed the penetrating bundle, which corresponds to the micro-CT image of the same, and later bifurcates into the left and right bundle branches (Fig 5e). The Purkinje fibers were visualized as red-stained structures along the endocardium layer, spread along the connective tissue (Fig 5f,g).

**Fig 5.**
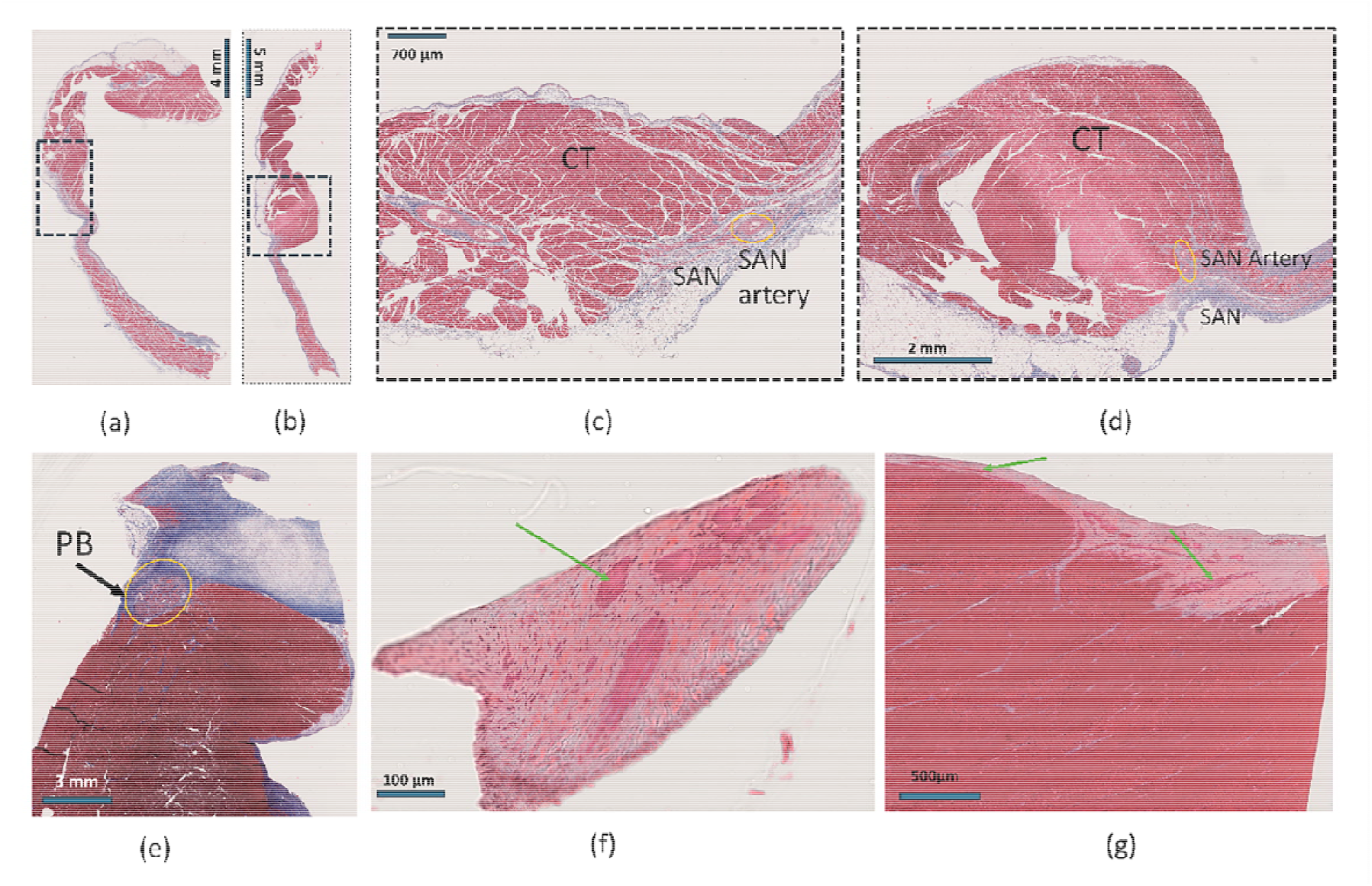
Histology images of (**a**) I_2_E stained SAN section (**b**) I_2_KI stained SAN section, (**c**) (d) Magnified images of marked regions in (a) and (b) respectively, showing the crista terminalis (CT), SAN region and SAN arteries (indicated by the blue arrows) (**e**) I_2_E stained AVN section showing the penetrating bundle (PB) (**f**) (**g**) Magnified images of the Purkinje fibers (PF-green arrows indicate the fibers running through the false tendons and endocardium respectively)

Sodium thiosulfate (STS) was found to be more effective than the conventional de-staining method that uses ethanol, which often took multiple weeks to remove iodine. This was observed for the de-staining of both I_2_E and I_2_KI samples. The contrast agents were cleared within 3 days of immersion in STS at 4°C (Fig 6).

**Fig 6.**
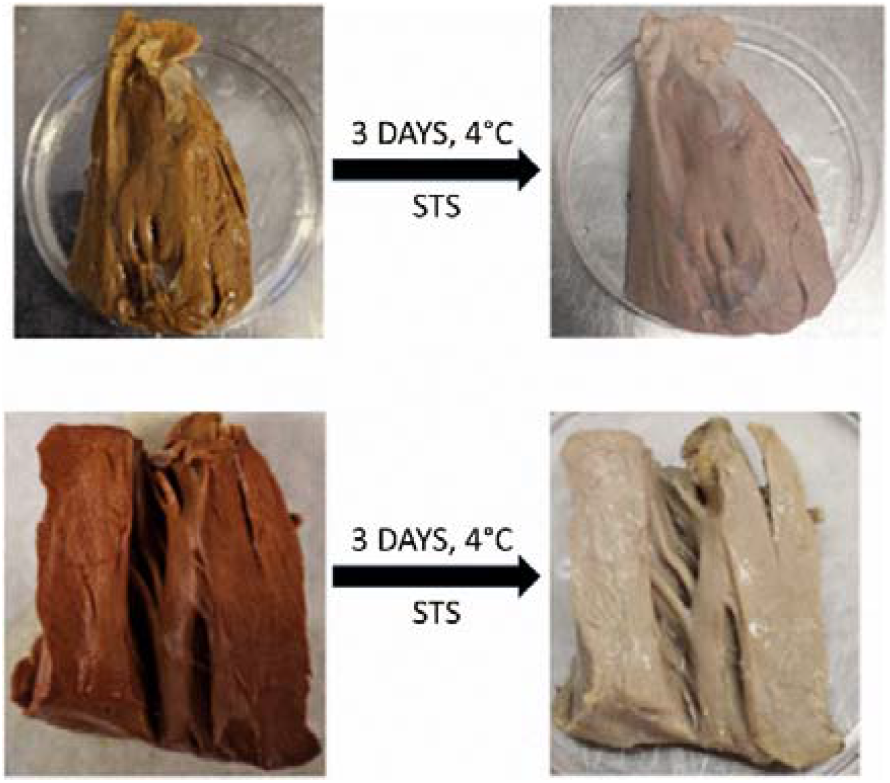
kI_2_KI removal using STS in the AVN section (top); I_2_E removal using STS in the apex section (bottom)

### Quantification of shrinkage

As expected, dehydration and iodine staining have together contributed to a progressive volumetric shrinkage in the I_2_E-stained sample. The use of ethanol induced a shrinkage of around 10% in the sample ahead of staining. The first day of staining resulted in a shrinkage of about 17% by volume compared to the native tissue volume. It was observed that there was almost a 46% decrease in volume after staining on day 5 when compared to the native volume. There was also a reduction of around 30% in volume between day 1 and day 5 of staining. The disintegration of the I_2_KI-stained sample resulted in a severe reduction in volume, which was observed within the initial two days, followed by a decrease in the rate of shrinkage thereafter. Overall, I_2_KI staining accounted for over 65% of the volumetric shrinkage during the entire staining period (Fig 7a,b).

**Fig 7.**
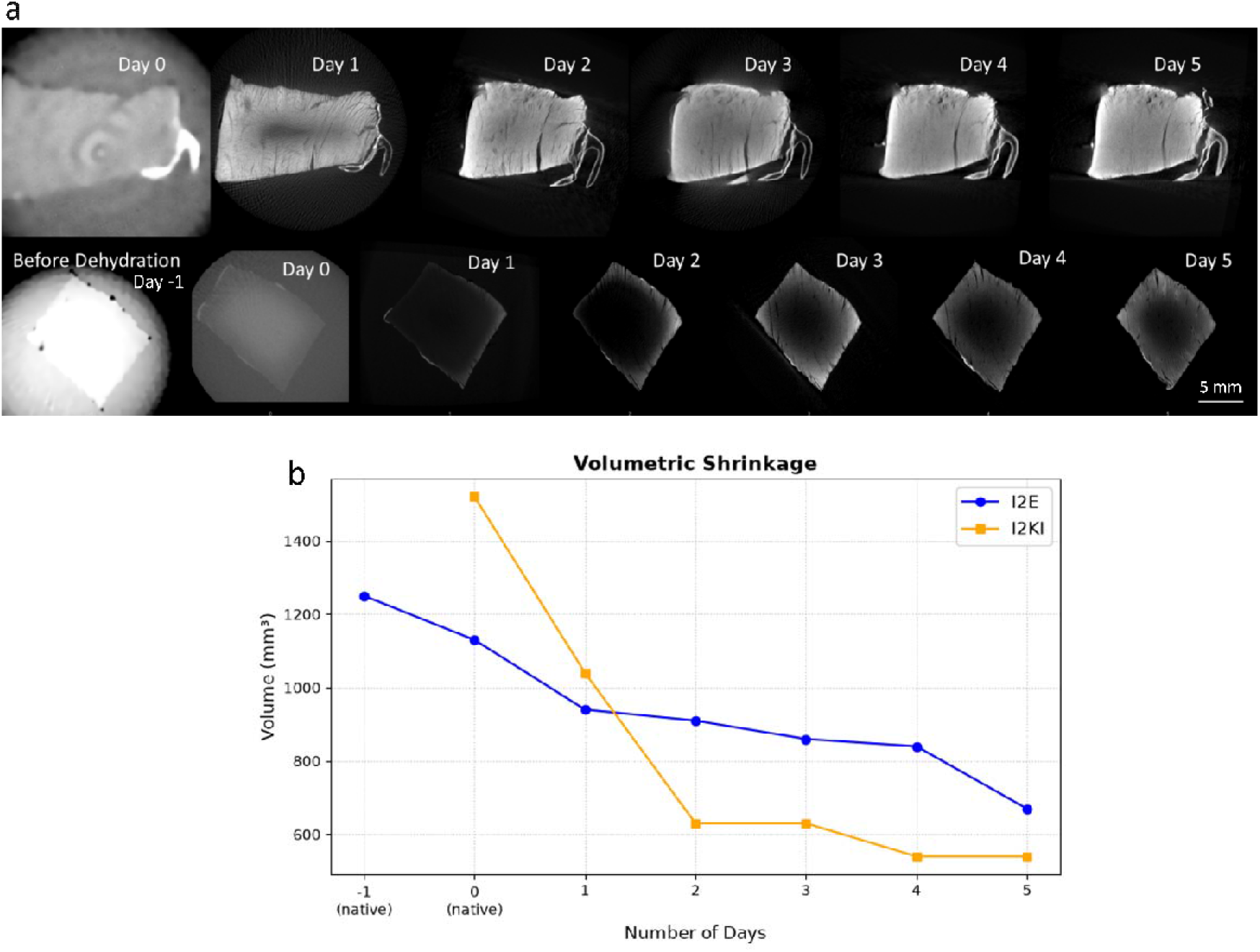
**(a)** Micro-CT images of I_2_KI-stained section (row 1) and I_2_E-stained section (row 2) showing the volumetric shrinkage over the staining period **(b)** Volumetric shrinkage for the I_2_E and I_2_KI-stained sample over the staining days. I_2_KI shows rapid disintegration within the initial staining days itself, while I_2_E exhibit slower, controlled shrinkage over time without disintegration.

### Image Quality Metrics

Fig 8 illustrates the statistical analyses on the contrast agents. The first panel demonstrates that the S-B Ratio is highly comparable between the two contrast agents. Both groups exhibit considerable intra-group variance and the associated p-value of 0.89 heavily exceeds the standard significance threshold (), indicating no statistically significant difference in the signal enhancement relative to the background.

**Fig 8.**
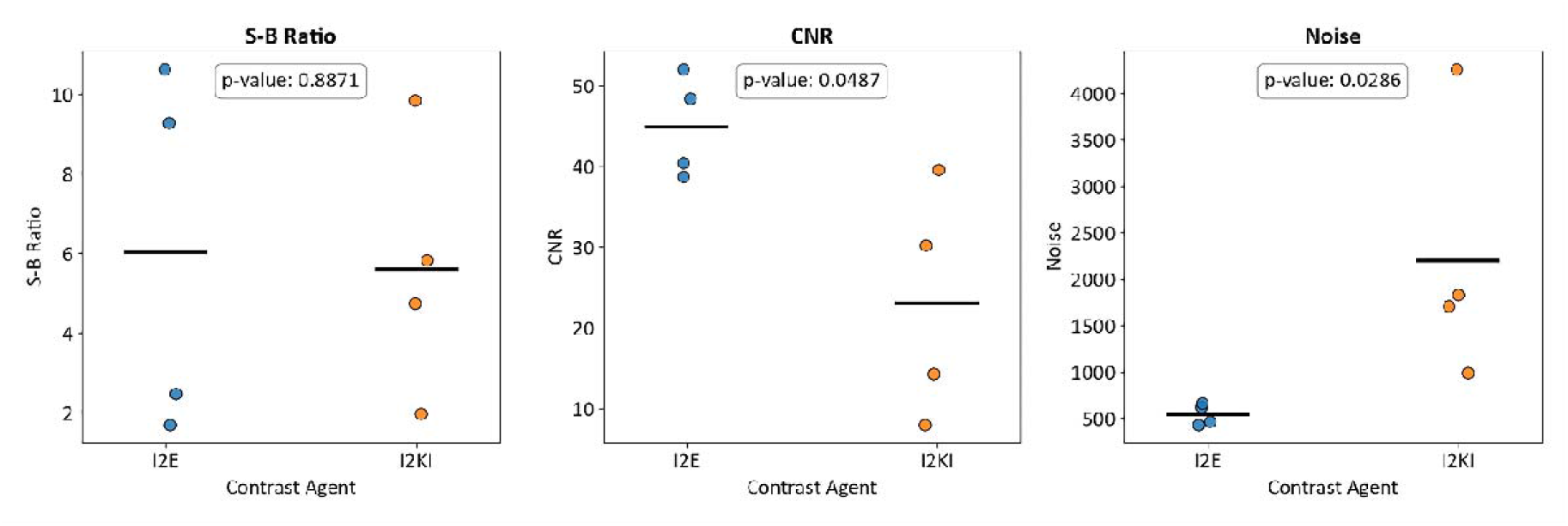
The strip plots display the Signal-to-Background (S-B) Ratio (left), Contrast-to-Noise Ratio (CNR) (middle), and Noise levels (right) for each agent. While there is no statistically significant difference in the S-B Ratio between the two contrast agents (), I_2_E demonstrates a significantly higher CNR () and significantly lower image noise () compared to I_2_KI.

When the CNR performance of both contrast agents was analyzed, I_2_E yields a substantially higher mean CNR compared to I_2_KI, which demonstrates a lower mean. The calculated p-value of 0.0487 indicates that this difference is statistically significant (). Consequently, I_2_E provides superior contrast differentiation relative to the inherent noise of the system.

The I_2_E cohort had persistently low noise levels with negligible variation across the samples, while the I_2_KI cohort shows a significantly elevated mean noise level coupled with considerable intra-group variability. The p-value of 0.0286 confirms a statistically significant reduction in noise when utilizing I_2_E compared to I_2_KI.

## Discussion

This research highlights the often-overlooked drawbacks of I_2_KI, including tissue disintegration, leaching, and staining-related artifacts. We propose I_2_E as an alternative staining agent and a protocol to assess the staining effectiveness of I_2_E and I_2_KI in distinguishing pertinent anatomical features while maintaining sample integrity for seamless subsequent processes.

The sample for I_2_E staining had to undergo a dehydration procedure before staining. Dehydration has been shown to increase cell membrane permeation and thereby allow the stain molecules to diffuse more easily into tissues. Furthermore, gradual dehydration aligns the tissue’s internal water content with the ethanol-based staining solution, facilitating a more chemically stable and uniform staining environment throughout the immersion phase [22][23]. However, prolonged storage of tissue in ethanol has also been reported to reduce and slow iodine uptake [24]. Although the exact mechanism of iodine interaction with molecular components is largely unknown and unexplored, triiodide ions in I_2_KI readily interact with glycogen and other soft-tissue components, increasing X-ray attenuation and signal intensity in soft tissue [25]. Previous studies show that iodine trimers bind to carbohydrates, such as glycogen, and to lipids through non-covalent interactions [26][27]. Myocardium has high glycogen content that strongly absorbs iodine, whereas ECM has modest absorption and lipids show solvent-dependent absorption. This could have resulted in the larger variation of signal intensity within samples of the same contrast agent. Furthermore, the ethanol dehydration in the case of I_2_E, dissolves lipids and proteins, thereby affecting iodine absorption.

I_2_KI staining, however, has produced artifacts, particularly background noise and streak artifacts. This could be due to the leaching of I_2_KI into the surrounding agarose, observed after micro-CT scanning and after melting the agarose. The leaching property of I_2_KI has also been reported in the studies by Aslanidi et al [28]. The higher noise and, thus the lower CNR could be due to the abovementioned issues with I_2_KI. The other major disadvantage of using I_2_KI as a staining agent is that it can disintegrate the sample, which prevents a smooth downstream process. Loss of such delicate tissue structures due to excessive breakage obscures various post-processing procedures with the samples, and it negates the advantage of micro-CT as a non-destructive imaging modality. Volumetric shrinkage analysis also clearly shows a sudden loss in volume within the first two days of I_2_KI staining. This deformation could be primarily attributed to the acidification of the staining solution over the staining period [17]. The sample had been largely broken by day 5 of staining. This makes sample preparation and staining more cumbersome, as fresh contrast agents must be prepared for subsequent staining. Furthermore, another study conducted by L. Costello et al. reveals that there is significant volumetric shrinkage during staining, especially at higher concentrations and longer durations. The study also demonstrates that high concentrations of I_2_KI (>5%) lead to oversaturation, resulting in poor contrast between muscle structures and making anatomical interpretation difficult [29]. The tissue shrinkage and disintegration were not caused by the I_2_/KI complex itself, but rather by the acidification of the solution. As iodine interacts with water and oxidizes biological tissue components, it generates free protons through hydrolysis [30]

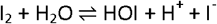

In the studies by Stephenson et al. using I_2_KI staining in an ex vivo setup, SAN was identified as the lightly stained region lying posterior to the crista terminalis (CT). Furthermore, in the AVN section, the penetrating bundle (PB) was identified as a circular mass that eventually divides into the left and right branches of the bundle of His[3]. These findings were consistent with those obtained when I_2_E was used in our study, and, moreover, SAN arteries and other vasculature were prominent, enabling their accurate segmentation. Thus, I_2_E can produce a better contrast in compartments enriched with lipids and proteins, such as adipose tissues and muscle constituents. However, I_2_KI staining did not give consistent results in identifying the CCS structure in terms of visual contrast. Moreover, the Purkinje fibers were the most challenging to identify due to their minute structure, and a low-resolution micro-CT scan was not sufficient for their imaging. The exact locations of the Purkinje fibers were identified with 2D images of the high-resolution scans and from previously published data that used different stains[31–33]. Even then, the high-resolution 3D images were found to be incapable of differentiating Purkinje fibers completely from false tendons and within ventricular myocardium.

Our study also corresponds with the findings of Reunamo et al. and Hopkins et al. in assessing the effectiveness of STS in clearing iodine staining[34,35]. Using STS as a clearing solution did not show any adverse effects in histological post-processing with Masson’s Trichrome staining. In the histological sections of the above-mentioned studies, the fibers were observed to be distributed in the endocardium, embedded within the collagen matrix. Additionally, we confirmed that the observed structures in 2D micro-CT images were Purkinje fibers based on their similarity in thickness with the histological images. The comparison between histological sections and micro-CT images was necessary to validate the anatomical accuracy and assess tissue integrity following staining protocols. Neither stain caused any adverse effects on the differentiation of anatomical features in histology. The histology images consistently showed the SAN and nodal arteries, penetrating bundle of the AVN that branches into the bundle of His and Purkinje fibers. However, the greater tissue disintegration exhibited by I_2_KI-stained sections is likely due to the acidification of the I_2_KI as mentioned above. This suggests a trade-off between enhanced signal intensity and preservation of fine tissue architecture, thereby questioning the robustness of the I_2_KI approach.

One of the limitations of this study is that the concentrations of contrast agents were not equal. For I_2_KI, the concentration was decided based on previous cardiac and other soft-tissue studies that suggested a concentration below 5% would give optimum results [19,25,36,37]. A 2.5% I_2_KI concentration was used based on the aforementioned cardiac studies. In the case of I_2_E, we found that a 1% concentration was sufficient to provide contrast for differentiating the CCS structures. Also, a larger cohort would have provided for a more robust result. Furthermore, the macroscopic tissue sizes varied, and the field of view had to be adjusted accordingly. There was thus a trade-off between resolution and field of view while imaging.

Future directions of this study include using combined micro-CT and histological analyses to deepen the understanding of cardiac pathologies, such as myocardial fibrosis, and to explore applications in drug testing. Additionally, the 3D imaging capabilities of micro-CT and segmentation of CCS structures offer opportunities to develop advanced computational models. These models could enhance research on rhythm disorders, support the design of cardiac devices, and, in turn, help optimize patient outcomes.

## Conclusion

In this work, we present iodine in ethanol as a suitable alternative for cardiac micro-CT research. To our knowledge, this is the first time I_2_E has been systematically evaluated against I_2_KI for imaging the cardiac conduction system in large mammalian (porcine) hearts. For cardiac imaging, the I_2_KI approach offers faster staining, with the trade-off of compromised sample integrity and structural preservation. Iodine in ethanol provides sufficient contrast for differentiating anatomical structures with minimal staining-related artifacts. Moreover, it supports a smooth downstream procedure, unlike I_2_KI, making it viable for further imaging and histological sectioning.

## Data Availability Declaration

The datasets generated and analyzed during this study are not publicly available but are available from the corresponding author upon request.

## Author Contribution Declaration

All authors contributed to the study conception and design. Formal analysis, investigation and writing was performed by Manu Pradeep, Conceptualization was done by Shuvashis Das Gupta, and Sami Kauppinen, Resource and material preparation was done by Yi Li. The experimental design and result interpretation was done by Mikko Finnilä and supervision was performed by Timo Liimatainen. All authors commented on previous versions of the manuscript. All authors read and approved the final manuscript.

## Statement and Declaration

The authors have no competing interests to declare that are relevant to the content of this article.

## Acknowledgement

We thank Tarja Huhta (senior laboratory technician, University of Oulu) for her assistance with histological tasks.

